# Enrich2: a statistical framework for analyzing deep mutational scanning data

**DOI:** 10.1101/075150

**Authors:** Alan F. Rubin, Nathan Lucas, Sandra M. Bajjalieh, Anthony T. Papenfuss, Terence P. Speed, Douglas M. Fowler

## Abstract

Measuring the functional consequences of protein variants can reveal how a protein works or help unlock the meaning of an individual’s genome. Deep mutational scanning is a widely used method for multiplex measurement of the functional consequences of protein variants. A major limitation of this method has been the lack of a common analysis framework. We developed a statistical model for estimating variant scores that can be applied to many experimental designs. Our method generates an error estimate for each score that captures both sampling error and consistency between replicates. We apply our model to one novel and five published datasets comprising 243,732 variants and demonstrate its superiority, particularly for removing noisy variants, detecting variants of small effect, and conducting hypothesis testing. We implemented our model in easy-to-use software, Enrich2, that can empower researchers analyzing deep mutational scanning data.

## Introduction

Exploring the relationship between sequence and function is fundamental to enhancing our understanding of biology, evolution, and genetically driven disease. Deep mutational scanning is a method that marries deep sequencing to selection amongst a large library of protein variants, measuring the functional consequences of hundreds of thousands of variants of a protein simultaneously. Deep mutational scanning has greatly enhanced our ability to probe the protein sequence-function relationship [1], and has become widely-used [2]. For example, deep mutational scanning has been applied to comprehensive interpretation of variants found in disease-related human genes [3], understanding protein evolution [4–8], and probing protein structure [9,10] with many additional possibilities on the horizon [2].

In a deep mutational scan, a library of protein variants is first introduced into a model system [11]. Model systems that have been used in deep mutational scanning include phage, bacteria, and yeast. A selection is applied for protein function or another molecular property of interest, altering the frequency of each variant according to its functional capacity. Selections can be growth-based or implement physical separation of variants into bins, as in phage display or flow sorting of cells. Next, the frequency of each variant in each time point or bin is determined by using deep sequencing to count the number of times each variant appears. Here, the variable region is either directly sequenced using a single- or paired-end strategy, or a short barcode that uniquely identifies each variant in the population is sequenced instead [11,12]. Analysis of the change in each variant’s frequency throughout the selection yields a score that estimates the variant’s effect. Scoring the performance of individual variants is distinct from a related class of methods that quantify tolerance for change at each position in a target protein [13]. Those approaches enable a different set of biological inferences that we do not seek to address here.

Fundamental gaps remain in our ability to use deep mutational scanning data to accurately measure the effect of each variant because practitioners lack a unifying statistical framework within which to interpret their results. Existing methods are diverse in terms of their scoring function, statistical approach, and generalizability. Two established implementations of deep mutational scanning scoring methods, Enrich [14] and EMPIRIC [15], calculate variant scores based on the ratio of variant frequencies before and after selection. This type of ratio-based scoring has been used to quantify the effect of noncoding changes in promoters as well [16]. However, while intuitive and easy to calculate, ratio-based scores are highly sensitive to sampling error when frequencies are low. For experimental designs that sample from more than two time points to improve the resolution of changes in frequency, ratio-based scoring is insufficient so a regression-based approach has been used instead [3,17–19]. Both ratio and regression analyses can incorporate corrections for wild type performance [7,14,15,18,20] or nonsense variants [15,17] at the expense of restricting the method to protein coding targets only.

The lack of a common standard for calculating scores makes comparison between studies difficult, and existing bespoke methods are not applicable to the diverse array of experimental designs currently being used. Furthermore, no existing method quantifies the uncertainty surrounding each score, which limits the utility of the data. For example, one of the most compelling applications of deep mutational scanning is to annotate variants found in human genomes with the goal of empowering variant interpretation [3], where estimation of the uncertainty associated with each measurement in a common framework is crucial. At best, current approaches employ *ad hoc* filtering of putative low-quality scores, often using manually determined read-depth cutoffs.

To address these limitations, we present Enrich2, an extensible and easy-to-use computational tool that implements a comprehensive statistical model for analyzing deep mutational scanning data. Enrich2 includes scoring methods applicable to deep mutational scans with any number of time points. Unlike existing methods, Enrich2 also estimates variant scores and standard errors that reflect both sampling error and consistency between replicates. We explore Enrich2 performance using novel and published deep mutational scanning data sets comprising 243,732 variants in five target proteins. We demonstrate that Enrich2’s scoring methods perform better than existing methods regardless of experimental design. Enrich2 facilitates superior removal of noisy variants and improved detection of variants of small effect, and enables statistically rigorous comparisons between variants. Enrich2 is platform-independent and includes a graphical interface designed to be accessible to experimental biologists with minimal bioinformatics experience.

## Results and discussion

### Overview of Enrich2 workflow

We distilled the common features of a deep mutational scan into a generalized workflow (Fig. 1). After the experiment, each FASTQ file is quality filtered and variants are counted. For directly sequenced libraries, this involves calling the variant for each read (see **Materials and methods**). For barcoded libraries, barcode counts are assigned to variants using an additional file that describes the many-to-one barcode-to-variant relationship. Next, the counts for each variant are normalized and a score is calculated that quantifies the change in frequency of each variant in each selection. Finally, each variant’s scores from replicate selections are combined into a single replicate score using a random-effects model. Variant standard errors are also calculated for each selection and replicate score, allowing the experimenter to remove noisy variants or perform hypothesis testing. Enrich2 is designed to enable users to implement other scoring functions, so long as they produce a score and a standard error. Thus, Enrich2 can serve as a framework for any counting-based enrichment/depletion experiment.

**Figure 1:**
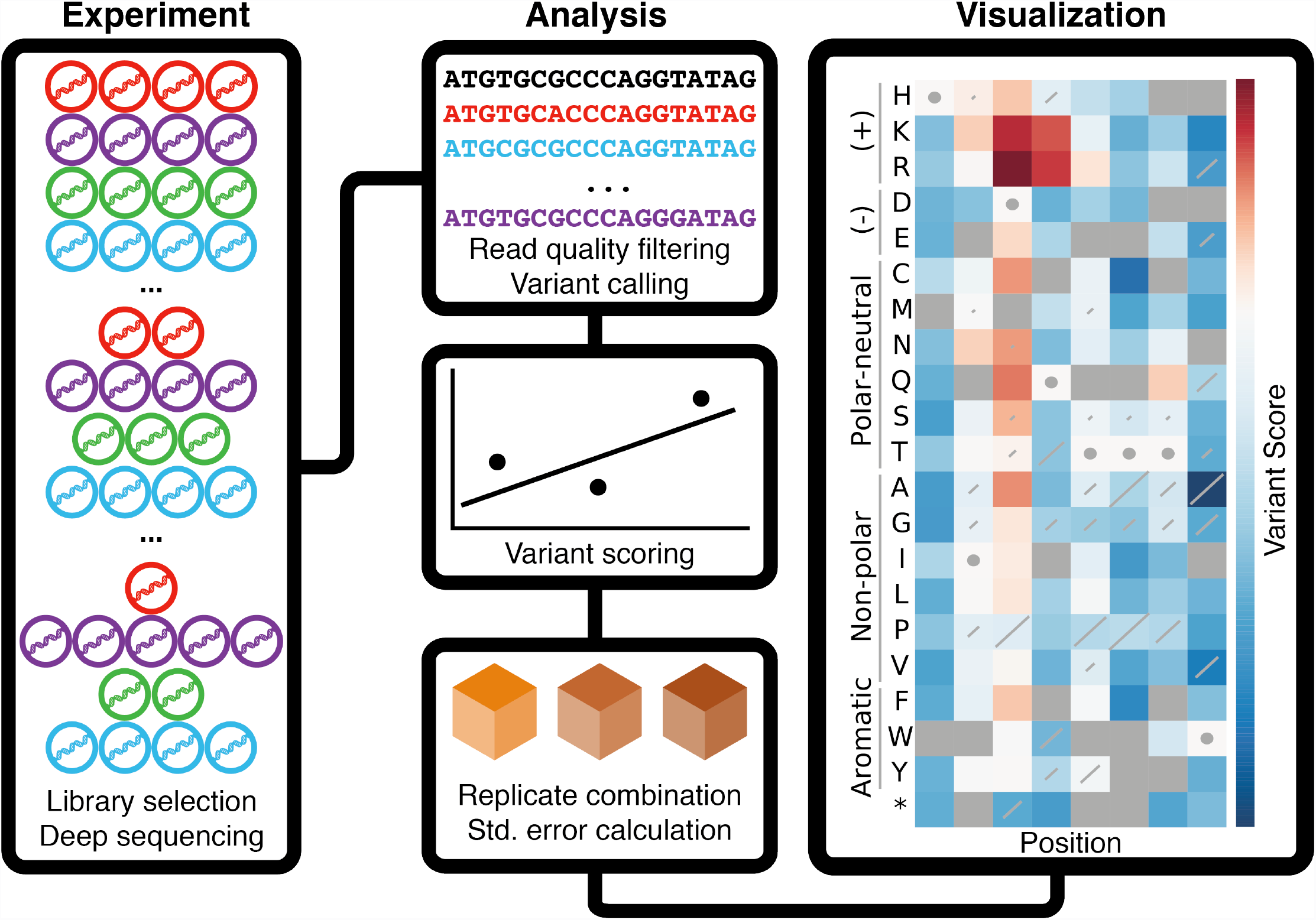
Deep mutational scanning and Enrich2. In a deep mutational scan, a library of protein variants is subjected to selection, which perturbs the frequency of variants. Samples of the library are collected before, during, and after selection and subjected to high-throughput sequencing (**left panel**). Enrich2 processes the high-throughput sequencing files generated from each sample. Sequencing reads are quality filtered, and variants are counted by comparing each read to the wild type sequence. Enrich2 estimates variant scores and standard errors using the variant counts, and combines these estimates for replicates (**middle panel**). Enrich2 displays the scores and standard errors as a sequence-function map. A sequence-function map of eight positions of the hYAP65 WW domain is shown (**right panel**). Cell color indicates the score for the single amino acid change (row) at the given position in the mutagenized region (column). Positive scores (in red) indicate better than wild type performance in the assay, and negative scores (in blue) indicate worse than wild type performance. Diagonal lines in each cell represent the standard error for the score, and are scaled such that the highest standard error on the plot covers the entire diagonal. Standard errors that are less than 2% of this maximum value are not plotted. Cells containing circles have the wild type amino acid at that position. Grey squares denote amino acid changes that were not measured in the assay.

### Scoring a single selection using linear regression

For experimental designs with three or more time points, Enrich2 calculates a score for each variant using weighted linear least squares regression. These time points can be variably spaced, as in samples from a yeast selection withdrawn at different times, or they can be uniformly spaced to represent rounds or bins, as in successive rounds of a phage selection. Each variant’s score is defined as the slope of the regression line. For each time point in the selection, including the input time point, we calculate a log ratio of the variant’s frequency relative to the wild type’s frequency in the same time point and regress these values on time. Regression weights are calculated for each variant in each time point based on the Poisson variance of the variant’s count (see **Materials and methods**). We estimate a standard error for each score using the weighted mean square of the residuals about the fitted line. We calculate *p*-values for each score using the *z*-distribution under the null hypothesis that the variant behaves like wild type (*i.e.* has slope of 0).

A problem with linear regression-based scoring is that the wild type frequency often changes non-linearly over time in an experiment-and selection-specific manner (Fig. 2). Existing linear model-based approaches subtract the wild type score from each variant’s score [3,17] ignoring this issue and potentially reducing score accuracy. A solution for this problem, normalizing each variant’s score to wild type at each time point, has been proposed but remained untested with real data [18]. We implemented per-time point normalization and compared variant standard errors calculated with and without wild type normalization for a total of 14 replicates in three different experiments: a phage selection for BRCA1 E3 ubiquitin ligase activity, a yeast two-hybrid selection for BRCA1-BARD1 binding, and a phage selection for E4B E3 ubiquitin ligase activity (Table 1). In all cases, wild type normalization resulted in significantly smaller variant standard errors (*p* ≈ 0, binomial test, **Table S1**). Variants that remain non-linear after normalization are poorly fit by our regression model and have high standard errors. Thus, they can easily be identified for further examination or removal. For experimental designs that do not have a wild type sequence, we normalize using the library size instead of the wild type count.

**Figure 2:**
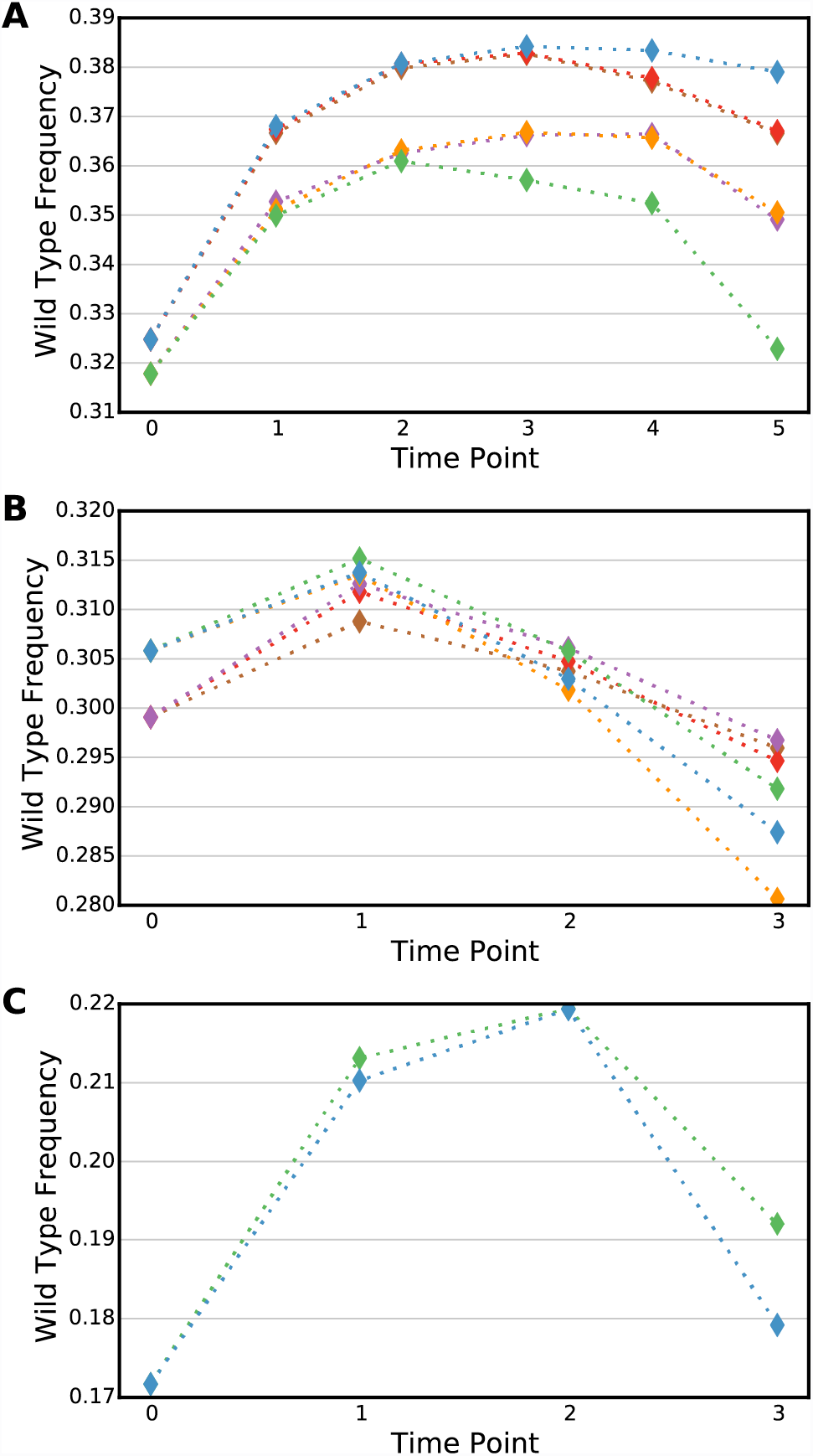
Wild type frequency can change non-linearly. The change in frequency of the wild type over the course of replicate selections is shown for (**A**) BRCA1 E3 ubiquitin ligase (**B**) BRCA1-BARD1 binding or (**C**) E4B E3 ubiquitin ligase. Each colored line represents a single replicate.

**Table 1:** Datasets analyzed with Enrich2

| Target | Assay | Replicates | Time Points | Scored Variants | Reads (millions) | Run Time (HH:MM) | Reference |
| --- | --- | --- | --- | --- | --- | --- | --- |
| BRCA1 | Phage display | 6 | 6 | 11,530 | 423 | 6:23 | [3] |
| BRCA1 | Yeast two-hybrid | 6 | 4 | 17,165 | 306 |  |  |
| E4B | Phage display | 2 | 4 | 158,939 | 67 | 2:27 | [21] |
| Neuraminidase | Growth in cell culture | 6* | 2 | 6,834 | 24 | 0:08 | [22] |
| C2 domain | Phage display | 3 | 3 | 1,081 | 48 | 1:17 | This work |
| WW domain | Phage display | 2 | 4 | 48,183 | 33 | 10:32 | [1] |

\* Three replicates each of two experimental conditions

Wild type non-linearity is not the only problem in scoring a typical selection. Each time point has a different number of reads per variant, and time points with low coverage are more affected by sampling error. An example of this issue is found in one of the replicate selections for BRCA1 E3 ubiquitin ligase activity (Fig. 3A). To address this problem, Enrich2 downweights time points in the regression with low counts per variant. Without weighted regression, the experimenter is forced to choose between three undesirable options: using the low coverage time point and adding noise to the measurements, removing the time point and complicating efforts to compare replicates, or spending time and resources to re-sequence the time point. Weighting avoids these undesirable options, achieving lower variant standard errors as compared to ordinary regression (Fig. 3B). To show that this effect is general and not a feature of the specific BRCA1 replicate we analyzed, we downsampled reads from a single time point in the E4B E3 ubiquitin ligase data set. We find that weighted regression reduces the mean standard error regardless of the fraction of reads removed (Fig. 3C, D). Finally, we show that weighted regression improves reproducibility between replicates in the BRCA1 E3 ubiquitin ligase data set even in the absence of any filtering (Fig 3E, F.).

**Figure 3:**
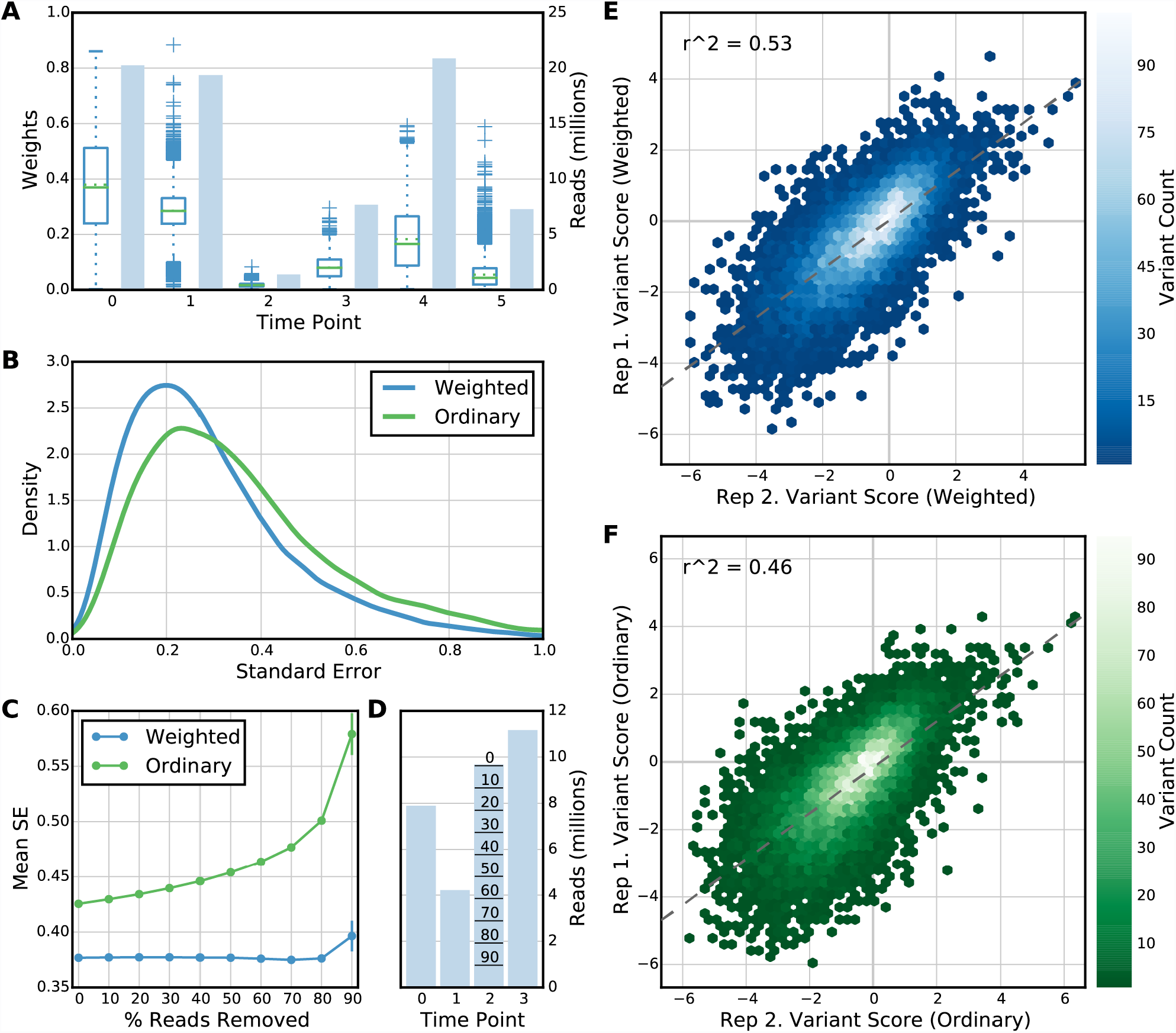
Weighted least squares regression reduces standard error and improves replicate correlation. (**A**) The number of reads (shaded blue bars) and the distribution of variant regression weights (boxplots, green line = median, dotted green line = mean) for each time point in a single BRCA1 E3 ubiquitin ligase selection is shown. Time points with fewer reads per variant are downweighted in the regression. The weights for later time points are lower on average because most variants decrease in frequency during the course of the selection. (**B**) A density plot of standard errors for all variants in the selection shown in panel A calculated using weighted least squares regression (blue line) or ordinary least squares regression (green line) is shown. The weighted least squares regression method returns lower standard errors using the same underlying data by minimizing the impact of sampling error in low read count time points. (**C**) The mean standard error of variants after randomly downsampling reads in a single time point in one of the E4B E3 ubiquitin ligase selections is shown. Mean standard errors for all variants at each read downsampling percentage were calculated using either weighted least squares (blue) or ordinary least squares (green) regression scoring. Error bars indicate the 95% confidence interval of five random downsampling trials at each percentage. (**D**) Read counts per time point in the selection described in panel C is shown. The lines on the bar for time point 2 correspond to the level of downsampling on the x-axis of panel C. (**E, F**) Plots of variant scores in two replicate selections from the BRCA1 E3 ubiquitin ligase data set are shown. Replicate agreement for scores calculated using the weighted least squares regression model (**E**) is higher than agreement for scores calculated using ordinary least squares regression (**F**). The dashed line shows the line of best fit for the replicate scores in each plot. Hex color indicates point density.

For experiments with only two sequenced populations or time points (e.g. “input” and “selected”), Enrich2 calculates the slope between the two time point log ratios, which is equivalent to frequently used ratio-based scoring methods [14,15,20]. Unlike previous implementations of ratio-based scoring, we provide standard error estimates for each score using Poisson assumptions (see **Materials and methods**).

### A random-effects model for scoring replicate selections

Deep mutational scans are affected by various sources of error in addition to sampling error. One way to deal with this problem is to perform replicates. Usually, each variant’s score is calculated by taking the mean across replicates, which ignores the distribution of replicate scores. Furthermore, if an error is calculated, it is derived only from the replicate scores’ distribution and ignores any error associated with each replicate score. One alternative is to combine replicate scores using a fixed-effect model [23]. We examined this approach for the BRCA1 E3 ubiquitin ligase data set (Fig. 4) and found that because variant scores can vary widely between replicates, this method dramatically underestimates the standard error of the combined variant score. We therefore implemented a random-effects model that estimates each variant’s score based on the distribution of that variant’s scores across all replicates. This random-effects model also produces a standard error estimate for each variant that captures selection-specific error as well as error arising from the distribution of replicate scores (see **Materials and methods**).

**Figure 4:**
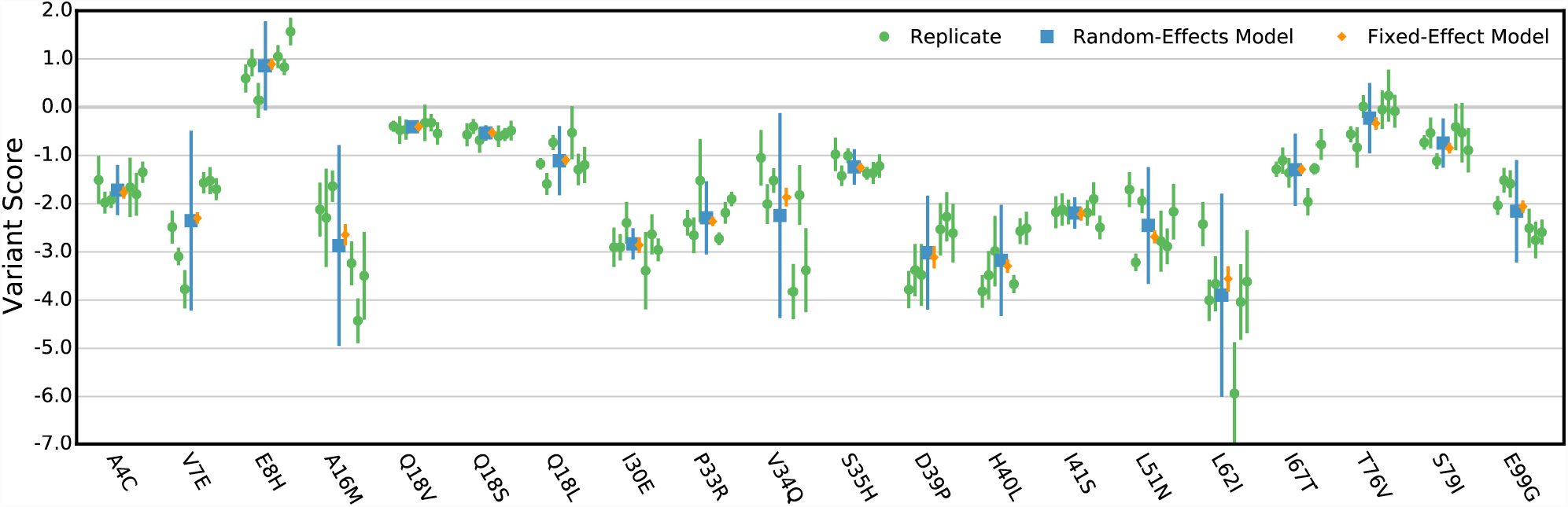
A random-effects model for scoring replicate selections. Variant scores for twenty randomly selected variants from the BRCA1 E3 ubiqutin ligase data set are shown. The replicate scores (green) were determined for each variant using Enrich2 weighted regression. Combined variant score estimates were determined using a fixed-effect model (orange) or the Enrich2 random-effects model (blue). In all cases, error bars show plus or minus two standard errors.

The random-effects model furnishes variant scores that are less sensitive to outlier replicates than a fixed-effect model (Fig. 4). Additionally, standard errors estimated by the random-effects model better reflect the distribution of replicate scores, providing a better basis for subsequent hypothesis testing. The same random-effects model can be used for experiments with any number of time points or replicates, or with any Enrich2 scoring function (Fig. S1). A key advantage of this approach is that error is quantified on a per-variant basis, unlike the usual approach of comparing replicate selections using pairwise correlation [3,13,17]. This allows experimenters to use replicate data to make inferences about individual variants, rather than simply as a quality control check for whole experiments.

### Standard error-based variant filtering

Per-variant standard error estimates enable the removal of variants with unreliable scores. This contrasts with previous filtering schemes, which employed an empirical cut-off for the minimum number of read counts for each variant in the input library or throughout the selection [1,3,21,22,24–29]. Read count cut-offs eliminate low-count variants that may be unreliably scored due to sampling error, but ignore other sources of noise and may introduce a bias against variants that become depleted after selection. Enrich2 retains low-count variants and enables the experimenter to determine which scores are reliable directly from the associated standard error.

To assess whether standard error-based filtering performs better than read count-based filtering, we analyzed data from a deep mutational scan of the C2 domain of Phospholipase A2 (Table 1). Here, a library of 84,252 phage-displayed C2 domain variants was selected for lipid binding over several rounds. This dataset was unanalyzable using previous methods due to the apparent extreme variability between replicate selections. We compared filtering based on three different parameters: Enrich2 variant standard error, read count in the input round, and total read count in all rounds of selection. To quantify filtering method performance, we took the top quartile of variants selected by each filtering method. Then, we calculated the pairwise Pearson correlation coefficient between variant scores for each possible pair of the three replicates in the C2 domain data set (Fig. 5, **Table S2**). We found that standard error-based filtering was the only method that recovered a replicable subset of variants from this data set. In fact, input count filtering selected a subset of variants whose scores were more poorly correlated than the unfiltered set. We performed a similar analysis on the much higher quality BRCA1 E3 ubiquitin ligase activity data set, and found that standard error filtering performed better (mean pairwise Pearson r^2^ = 0.89) than input library count (r^2^ = 0.84) or total count (r^2^ = 0.84) (**Table S3**).

**Figure 5:**
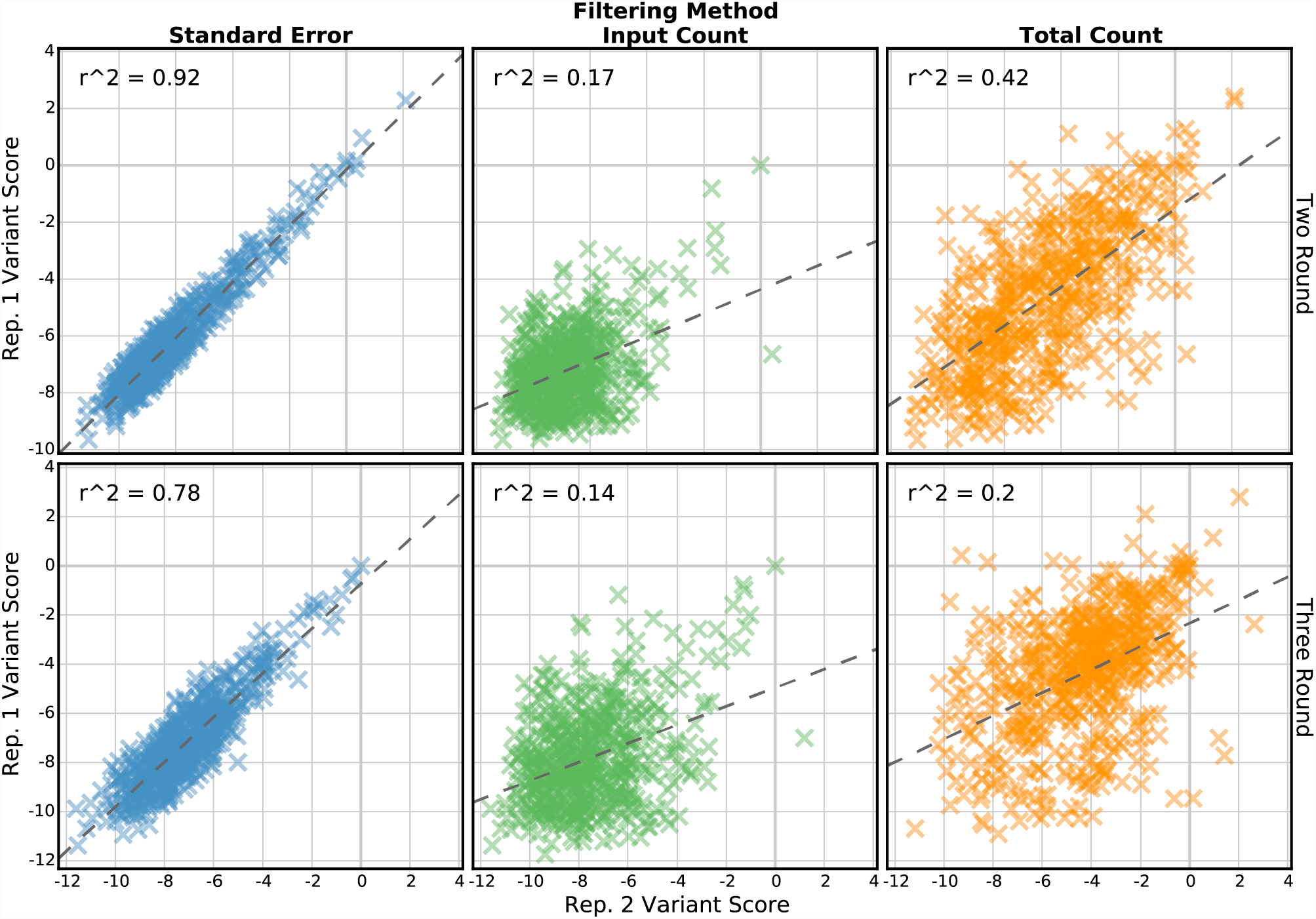
Standard error-based filtering improves replicate correlation. Variant scores from two replicates of the C2 domain data set are shown. Each panel plots the top quartile of variants selected by standard error (**left column**), input library count (**middle column**), or total count in all libraries (**right column**). Scores and standard errors are calculated using only the input and final round of selection (**top row**) or using all three rounds (**bottom row**). The dashed line is the best linear fit, and the Pearson correlation coefficient is shown.

To further demonstrate the utility of standard error-based filtering, we re-analyzed a deep mutational scan of the influenza virus neuraminidase gene (Table 1). In this experiment, 22 neuraminidase variants were individually validated and used to assess the quality of the deep mutational scanning data. Of these individually validated variants, four had large variant score standard errors as determined by Enrich2’s random-effects model (Fig. 6A, Fig. S1, **Table S4**). Removing these high-standard-error variants improved the correlation between the deep mutational scanning scores and individual validation scores from Pearson r^2^ = 0.81 to r^2^ = 0.87. Removal of these scores also improved the correlation when variant scores were calculated as originally described in the study (Pearson r^2^ = 0.80 vs. r^2^ = 0.84) (Fig. S2) [22]. This suggests that scores of variants with low Enrich2 standard errors are more likely to reflect the results of gold standard validation experiments, and supports the use of standard error-based filtering for selecting candidate variants for follow up studies.

**Figure 6:**
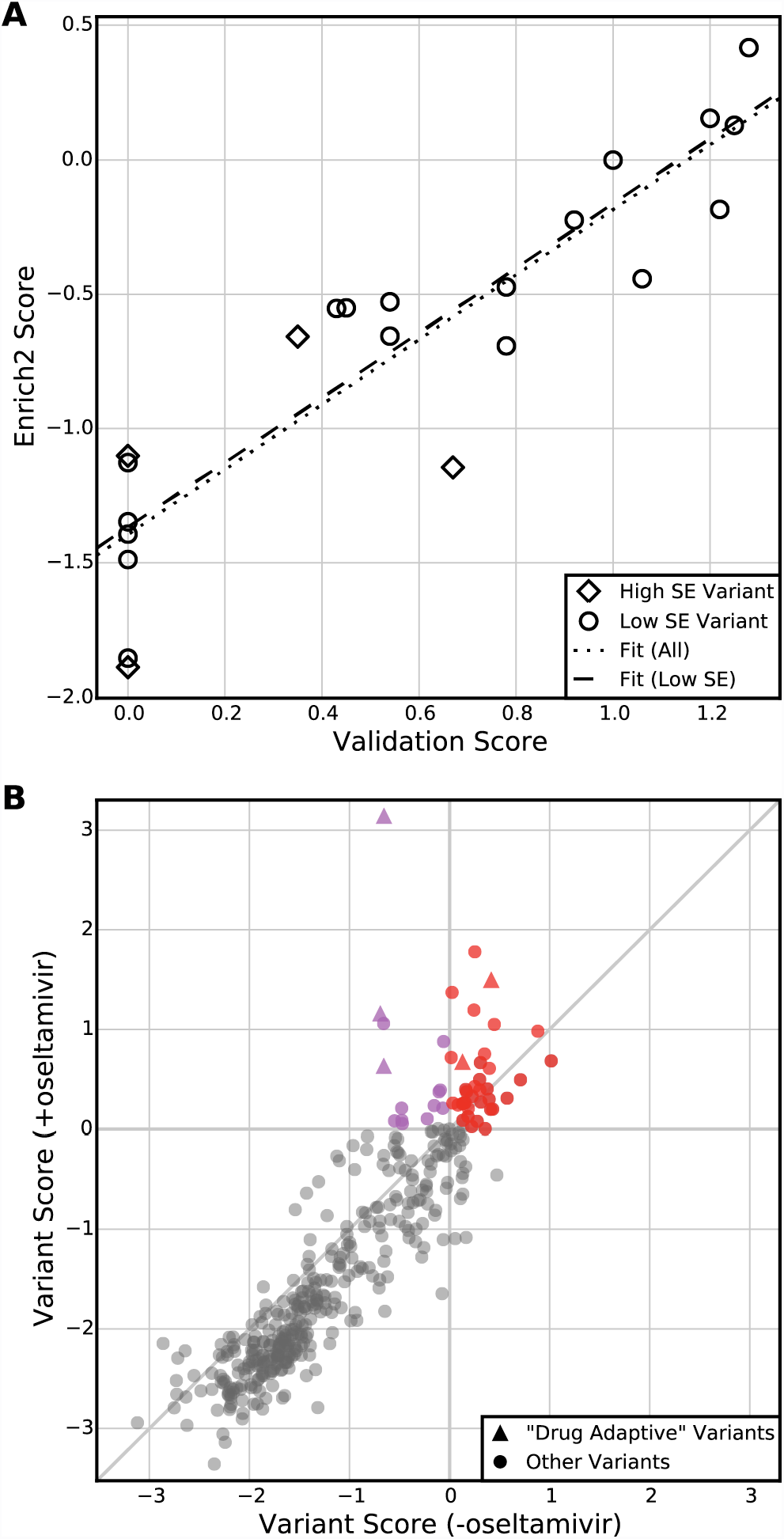
Standard errors enable hypothesis testing. (**A**) Enrich2 variant scores are plotted against single-variant growth assay scores for the 22 individually validated variants of the neuraminidase data set. Four (18%) of these variants have Enrich2 standard errors larger than the median standard error. The dotted line shows the best linear fit for all variants, and the dashed line shows the best linear fit for variants with standard errors less than the median. (**B**) Enrich2 variant scores are plotted for selections performed in the presence or absence of the small molecule inhibitor oseltamivir. Colored points indicate variants that significantly outperformed wild type in the drug’s presence. Red points also scored significantly higher than wild type in the drug’s absence. Triangles indicate the five “drug adaptive” mutations identified originally [22].

### Standard error-based hypothesis testing

An important challenge in analyzing deep mutational scanning data is determining whether a variant behaves differently from wild type or differently under altered conditions. Enrich2 standard errors empower experimenters to perform statistical tests for such differences. By default, Enrich2 calculates raw *p*-values for each score under the null hypothesis that the variant’s score is indistinguishable from wild type using a *z*-test. This allows the user to discriminate between variants with extreme scores due to sampling error or other noise from those that are confidently estimated to be different from wild type. We note that Enrich2 provides raw *p*-values, and users should correct for multiple testing using their preferred method.

We can also use a *z*-test to determine whether variants have different functional consequences under altered experimental conditions. For example, deep mutational scans of the neuraminidase gene were conducted in the presence and absence of the small molecule neuraminidase inhibitor oseltamivir (Table 1). The original study identified five “drug adaptive” variants, defined as those that outperformed wild type in the presence of oseltamivir [22]. These five drug adaptive variants included three known oseltamivir resistant variants. In our reanalysis, we identified 50 drug adaptive variants including all five variants found in the original study (Fig. 6B, **Table S5**). 36 of these 50 drug adaptive variants also had a significantly higher score than wild type in the absence of the inhibitor, and therefore might be more likely to occur in natural virus populations. Our results agree broadly with the original analysis, but by using Enrich2 to calculate scores and standard errors for variants across replicates, we were able to identify additional variants that may be of biological interest.

## Conclusions

We developed a unifying statistical framework for analyzing deep mutational scanning data that is applicable to most experimental designs. We showed that our statistical method is superior to existing methods for removing noisy variants and detecting variants of small effect, resulting in the identification of greater numbers of biologically interesting variants and enabling researchers to extract more from their datasets. We implemented our method in Enrich2, a computationally efficient graphical software package intended to improve access to deep mutational scanning for labs without data analysis experience. Enrich2 is extensible, so users can implement and easily share new scoring functions as new deep mutational scanning experimental designs are developed.

Enrich2 builds upon previous approaches to regression-based scoring which we improved in two ways. First, per-time-point wild type normalization helps reduce the effects of non-linear behavior under the assumption that many sources of non-linearity affect most variants similarly. Second, weighting each regression time point based on variant counts helps alleviate sampling error. In addition to these improvements, Enrich2 combines replicate selections into a single set of variant scores with standard errors to help identify variants that behave consistently in a given assay. Though variant score precision does not guarantee accuracy, we showed that removing variants with high standard errors from the neuraminidase data set did improve the correlation between deep mutational scanning results and gold-standard measurements.

Enrich2 standard errors can also be used to conduct hypothesis tests comparing variants within a single experimental condition or between multiple conditions. When comparing variants between conditions, we assume that the distribution of scores between conditions is roughly similar, but this assumption does not hold in all cases. For example, the shape of the score distribution is a function of the strength of the selective pressure applied [7,8]. Thus, Enrich2 standard errors should be used with caution when comparing variants between differing selections unless the variant scores are similarly distributed. A general method for normalizing scores to facilitate comparisons across different conditions or selection pressures remains an important open question.

The use of deep mutational scanning is expanding rapidly, and better tools for analysis will help it flourish. As with other widely-used high throughput experimental methods, a robustly implemented common statistical framework reduces barriers to entry, ensures data quality, and enables comparative analyses. We suggest that Enrich2 can help deep mutational scanning continue to grow by providing a foundation for meeting these challenges and facilitating further exploration and collaboration.

## Materials and methods

### Variant calling and sequence read handling

Enrich2 implements alignment-free variant calling [14]. Variant sequences are expected to have the same length and start point as the user-supplied wild type sequence, which allows Enrich2 to compare each variant to the wild type sequence in a computationally efficient manner. In addition to this alignment-free mode, an implementation of the Needleman-Wunsch global alignment algorithm [30] is included that will call insertion and deletion events. Enrich2 supports overlapping paired-end reads and single-end reads for direct variant sequencing, as well as barcode sequencing for barcode tagged variants,

### Calculating enrichment scores

For selections with at least three time points, we define *T*, which includes all time points, and *T*′, which includes all time points except the input (*t*_*0*_). The frequency of a variant (or barcode) *v* in time point *t* is the count of the variant in the time point (*c*_*v,t*_) divided by the number of reads sequenced in the time point (*N*_*t*_).

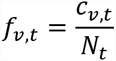

The change in frequency for a variant *v* in a non-input time point *t* ∈ *T*′ is the ratio of frequencies for *t* and the input.

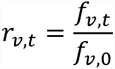

Instead of using this raw change in variant frequency, we divide each variant's ratio by the wild type (*wt*) variant's ratio.

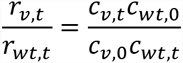

Because the library size terms (*N*_*t*_ and *N*_*0*_) in the frequencies cancel out, the ratio of ratios is not dependent on other non-wild type variants in the selection. In practice, we add 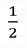 to each count to assist with very small counts [31] and take the natural log of this ratio of ratios.

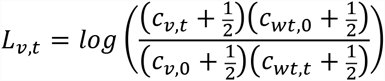

This equation can be rewritten as

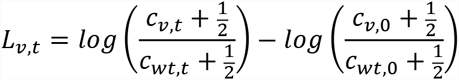

If we were to regress *L*_*v,t*_ on *t* ∈ *T*′, we note that the second term is shared between all the time points and therefore only affects the intercept of the regression line. We do not use the intercept in the score, so instead we regress on *M*_*v,t*_ and use all values of *t* ∈ *T*.

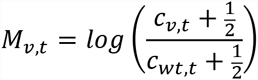

The score is defined as the slope of the regression line, 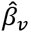. In practice, we regress on 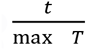 to facilitate comparisons between selections with different magnitudes of time points (e.g., 0/1/2/3 rounds vs. 0/24/48/72 hours).

To account for unequal information content across time points with variable sequencing coverage, we perform weighted linear least squares regression [32]. The regression weight for *M*_*v,t*_ is *V*_*v,t*_^−1^, where *V*_*v,t*_ is the variance of *M*_*v,t*_ based on Poisson assumptions [31] and is approximately

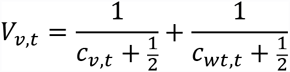

For selections with only two time points (e.g. input and selected), we use the slope of the line connecting the two points as the score. This is equivalent to the wild type adjusted log ratio (*L*_*v*_) derived similarly to *L*_*v,t*_ above.

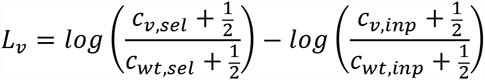

As there is no residual error about the fitted line, we must use a different method to estimate the standard error. We calculate a standard error (*SE*_*v*_) for the enrichment score *L*_*v*_ under Poisson assumptions [20,31].

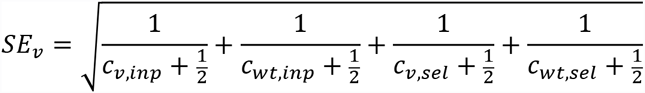

For experiments with no wild type sequence, scores can be calculated using the filtered library size for each time point *t*, which is defined as the sum of counts at time *t* for variants that are present in all time points.

### Random-effects model for combining replicate scores

To account for replicate heterogeneity, we use a simple meta-analysis model with a single random effect to combine scores from each of the *n* replicate selections into a single score for each variant. Each variant’s score is calculated independently. Enrich2 computes the restricted maximum likelihood estimates for the variant score 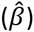 and standard error 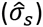 using Fisher scoring iterations [33]. Given the replicate scores 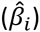 and estimated standard errors 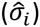 where *i = 1, 2,…, n,* the estimate for 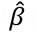 at each iteration is the weighted average:

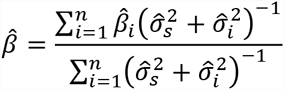

The starting value for 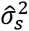 at the first iteration is:

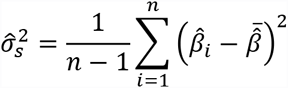

Enrich2 calculates the following fixed-point solution for 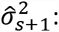

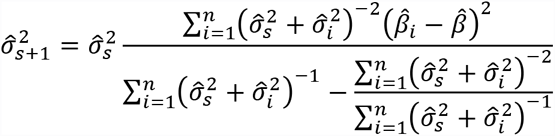

Because it is more computationally efficient to perform a fixed number of iterations for all variant scores in parallel than to test for convergence of each variant, Enrich2 performs 50 Fisher scoring iterations. In practice, this is more than sufficient for 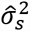 to converge. We record the difference 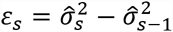 for the final iteration and identify any variants with high values for *ε*_*s*_ as variants that failed to converge. No such variants were encountered in the analyses detailed here.

### Deep mutational scan of Phospholipase A2

A region proximal to both lipid binding sites of the C2 domain of Phospholipase A2 (PLA2) was targeted for deep mutational scanning. Positions 94–97 of the C2 domain of mouse PLA2-alpha (ANYV) were fully randomized using a doped synthetic oligonucleotide. The library of C2 subdomains containing mutations was cloned into the AvrII and PpuMI sites of wild-type C2 domain in pGEM. The library was subcloned into phage arms and expressed on the surface of bacteriophage using the T7 phage display system according to the manufacturer’s instructions (Novagen T7Select 10–3b). The library was amplified in BLT5403 *E. coli* and variants were selected for their ability to bind to a lipid mixture containing ceramide 1-phosphate (C1P) [34]. The mouse PLA2-alpha cDNA was a generous gift from Michael Gelb, University of Washington. NiSepaharose Excel, capacity 10 mg/mL, was purchased from GE. Other reagents were purchased from Thermo-Fisher.

To select for C1P binding, lipid nanodiscs were developed as a bilayer affinity matrix. The His6-tagged membrane scaffold protein MSP1D1 [35] was expressed in BL21 *E. coli* from a pET28a plasmid and purified on nickel resin, then used to generate lipid nanodiscs comprised of 30 mol% phosphatidylcholine, 20 mol% phosphatidylserine, 40 mol% phosphatidylethanolamine, and 10 mol% C1P [36]. To separate nanodiscs from large lipid aggregates and free protein, the mixture was subjected to gel filtration using a Superose 6 10/300 GL column (Pharmacia) and the major peak following the void volume was collected. To generate the affinity resin, 70 µg of nanodiscs (quantified by protein content) was incubated overnight at 4 °C with 10 µl nickel resin in 20 mM Tris pH 7.5 and 100 mM NaCl. The resin was washed twice in the same solution and used in phage binding reactions.

Phage expressing the C2 domain variant library were titered and diluted to a concentration of 5 x 10^9^ pfu/mL in 20 mM Tris pH 7.5 and 100 mM NaCl, then incubated with lipid nanodisc affinity resin plus 10 µM calcium in a final volume of 350 µL. After a two-hour incubation at 4°C, the resin was washed four times in 1 mL of the incubation buffer containing 20 mM imidazole. Phage bound to nanodiscs were eluted with 20 mM Tris pH 7.5 containing 500mM imidazole. Phage from the elution were titered, amplified, and subjected to additional rounds of selection. Three replicate selections were performed on different days using the same input phage library.

Sequencing libraries were prepared by PCR amplifying the variable region using primers that append Illumina cluster generating and index sequences (**Table S6**) before sequencing using the Illumina NextSeq platform with a NextSeq high output kit (75 cycles, FC404-1005). Reads were demultiplexed using bcl2fastq v2.17 (Illumina) with the arguments *bcl2fastq --with-failed-reads --create-fastq-for-index-reads --no-lane-splitting --minimum-trimmed-read-length 0 --mask-short-adapter-reads 0*. Quality was assessed using FastQC v0.11.3 [37]. Demultiplexed reads are available in the NCBI Sequence Read Archive, BioProject Accession PRJNA344387.

### Neuraminidase data analysis

Raw reads were demultiplexed using a custom script based on three-nucleotide barcodes provided by the original authors [22]. The reads were analyzed in Enrich2 v1.0.0 as ten experimental conditions: five non-overlapping 30-base regions of the neuraminidase gene in either the presence or absence of oseltamivir. Reads were required to have a minimum quality score of 23 at all positions and contain no N’s. The five mutagenized regions were scored independently and then merged to create a single set of variant scores for each treatment. The original study excluded variants that were not intentionally generated by the mutagenesis approach employed, but we considered all single-amino acid variants that were present in all replicates and passed our quality filters. The *p*-values for comparing variant scores to wild type in each treatment and comparing variant scores between treatments were calculated using a z-test. All three sets of *p*-values were jointly corrected for multiple testing using the q-value package in R [38], and variants with a q-value of less than 0.05 were reported as significant.

### Analysis of other datasets

For previously published datasets, raw sequence files in FASTQ format were obtained from the respective authors. Datasets (Table 1) were analyzed independently using Enrich2 v1.0.0. The BRCA1 dataset was analyzed in a single run with separate experimental conditions for the yeast two-hybrid and phage display assays. For all datasets except neuraminidase, reads were required to have a minimum quality score of 20 at all positions and contain no N’s.

For the WW domain sequence function map (Fig. 1), scores and standard errors were calculated using weighted least squares linear regression in two technical replicates, and the replicates were combined using the random-effects model as described.

Estimated scores and standard errors for the fixed-effect model (Fig. 4), were calculated as described in [23].

### Enrich2 software implementation

Enrich2 is implemented in Python 2.7 and requires common dependencies for scientific Python. The graphical user interface is implemented using Tkinter. A deep mutational scanning experiment is represented as a tree of objects with four levels: experiment, condition, selection, and sequencing library. Each object’s data and metadata are stored in a single HDF5 file, including intermediate values calculated during analysis.

Enrich2 is designed to be run locally on a laptop computer and does not require a high performance computing environment. Most analyses can be run overnight (Table 1). Run times in Table 1 were measured using a MacBook Pro Retina with 2.8 GHz Intel Core i7 processor and 16GB of RAM.

The software is freely available from https://github.com/FowlerLab/Enrich2/ under the GPLv3 license. An example dataset and configuration file can be downloaded from https://github.com/FowlerLab/Enrich2-Example/. Online documentation, including implementation details, is located at http://enrich2.readthedocs.io/.

## Authors’ contributions

A.F.R., T.P.S., and D.M.F. developed the statistical methods. A.F.R. wrote the Enrich2 software and performed the data analysis. N.L., S.M.B., and D.M.F. designed and performed the C2 domain deep mutational scan. A.T.P. reviewed the codebase. A.F.R. and D.M.F. wrote the paper.

## Acknowledgements

We thank Stanley Fields for support, advice, and comments. We thank Lea Starita, Vanessa Gray, and Matt Rich for beta testing. We thank Bernie Pope and Matthew Wakefield for software engineering and algorithm advice. This work was supported by the National Institute of General Medical Sciences (1R01GM109110 and 5R24GM115277 to D.M.F. and P41GM103533 to Stanley Fields); the National Institute of Biomedical Imaging and Bioengineering (5R21EB020277 to S.M.B. and D.M.F.); and the National Health and Medical Research Council of Australia (Program Grant 1054618 to A.T.P. and T.P.S.).

---

**Figure S1:**
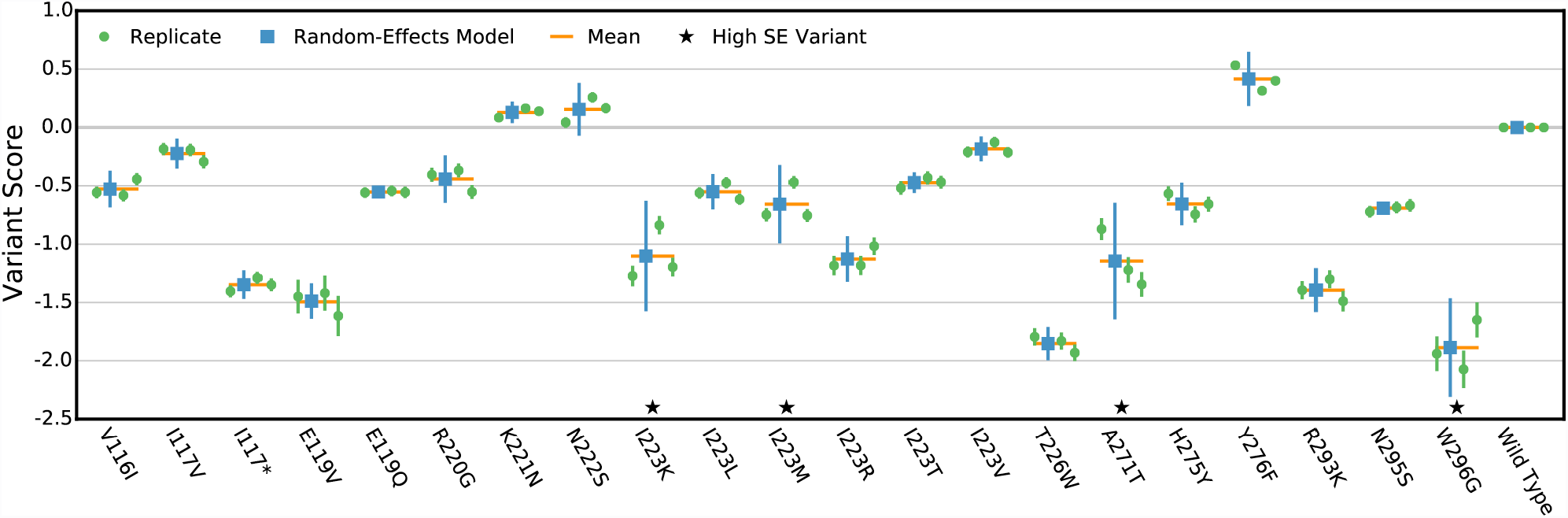
Random-effects model performance for individually validated neuraminidase variants. Variant scores for 22 individually validated variants from the neuraminidase data set are shown. The variant scores for each replicate (green) are plotted along with the mean variant score (orange line) and the combined variant score from the random-effects model (blue). Error bars on the replicate and random-effects model scores show plus or minus two standard errors. Variants with high standard errors (greater than 50^th^ percentile of scored variants with a single amino acid change) are marked with a star.

**Figure S2:**
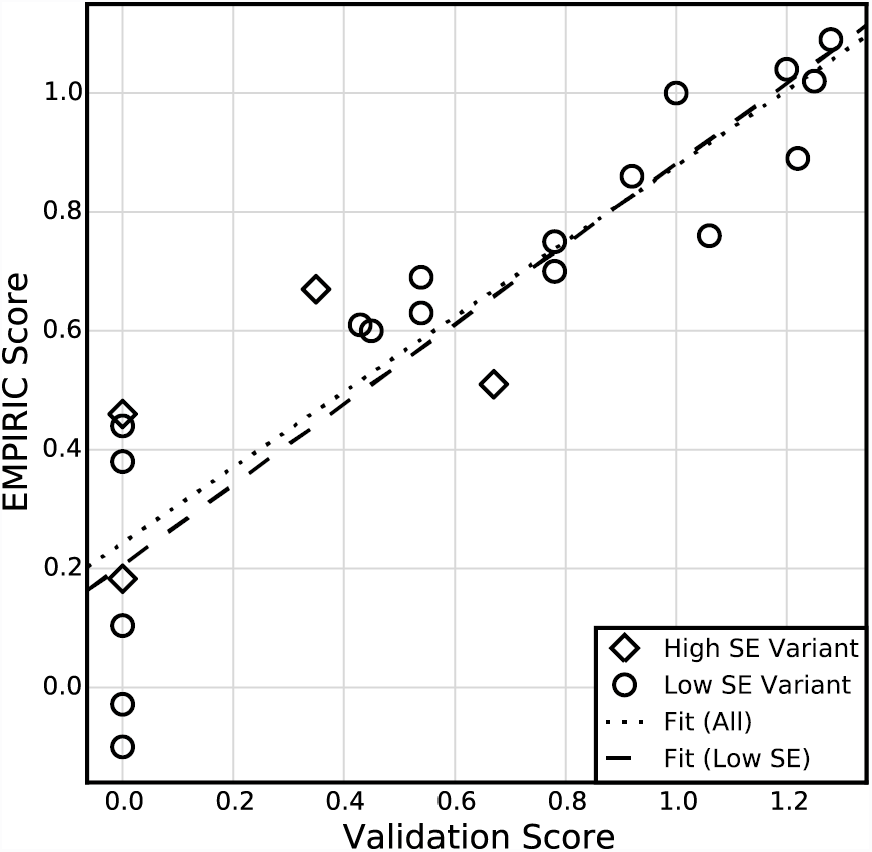
Standard errors enable hypothesis testing. EMPIRIC scores are plotted against single-variant growth assay scores for the individually validated variants of the neuraminidase data set. Four (18%) of these variants have Enrich2 standard errors larger than the median standard error. The dotted line shows the best linear fit for all variants, and the dashed line shows the best linear fit for variants with standard errors less than the median.

